# A porcine ex vivo model of pigmentary glaucoma

**DOI:** 10.1101/118448

**Authors:** Yalong Dang, Susannah Waxman, Chao Wang, Ralista T. Loewen, Ming Sun, Nils A. Loewen

## Abstract

Pigment dispersion syndrome can lead to pigmentary glaucoma (PG), a poorly understood condition of younger, myopic eyes with fluctuating, high intraocular pressure (IOP). The absence of a model similar in size and behavior to human eyes has made it difficult to investigate its pathogenesis. Here, we present a porcine ex vivo model that recreates the features of PG including intraocular hypertension, pigment accumulation in the trabecular meshwork and relative failure of phagocytosis. In *in vitro* monolayer cultures as well as in *ex vivo* eye perfusion cultures, we found that the trabecular meshwork (TM) cells that regulate outflow, form actin stress fibers and have a decreased phagocytosis. Gene expression microarray and pathway analysis indicated key roles of RhoA in regulating the TM cytoskeleton, motility, and phagocytosis thereby providing new targets for PG therapy.

## Introduction

Pigmentary glaucoma (PG) is a secondary open angle glaucoma in myopic eyes that affects patients in their 30s to 40s^1^. Individuals often experience a high, fluctuating intraocular pressure (IOP) that is more resistant to nonsurgical treatment than in primary open angle glaucoma^1, 2^. In addition to a baseline dispersion of pigment, physical activity^3, 4^ or eye movements can trigger pigment showers in some subjects, but patients often remain asymptomatic, making this condition particularly vexing. First described by Sugar and Barbour in 1949^5^, the clinical hallmark of pigment release is readily apparent with transillumination of the mid-peripheral iris (Fig. 1), deposition of pigment on the corneal endothelium (Krukenberg spindle) and in the trabecular meshwork (TM)^6^ yet the pathogenesis remains poorly understood. Pigment dispersion seems to be caused by mutations or variants of more than one gene. Although a susceptibility locus was mapped to chromosome 7q35–q36, a candidate gene remains yet to be identified^7^.

**Figure 1:**
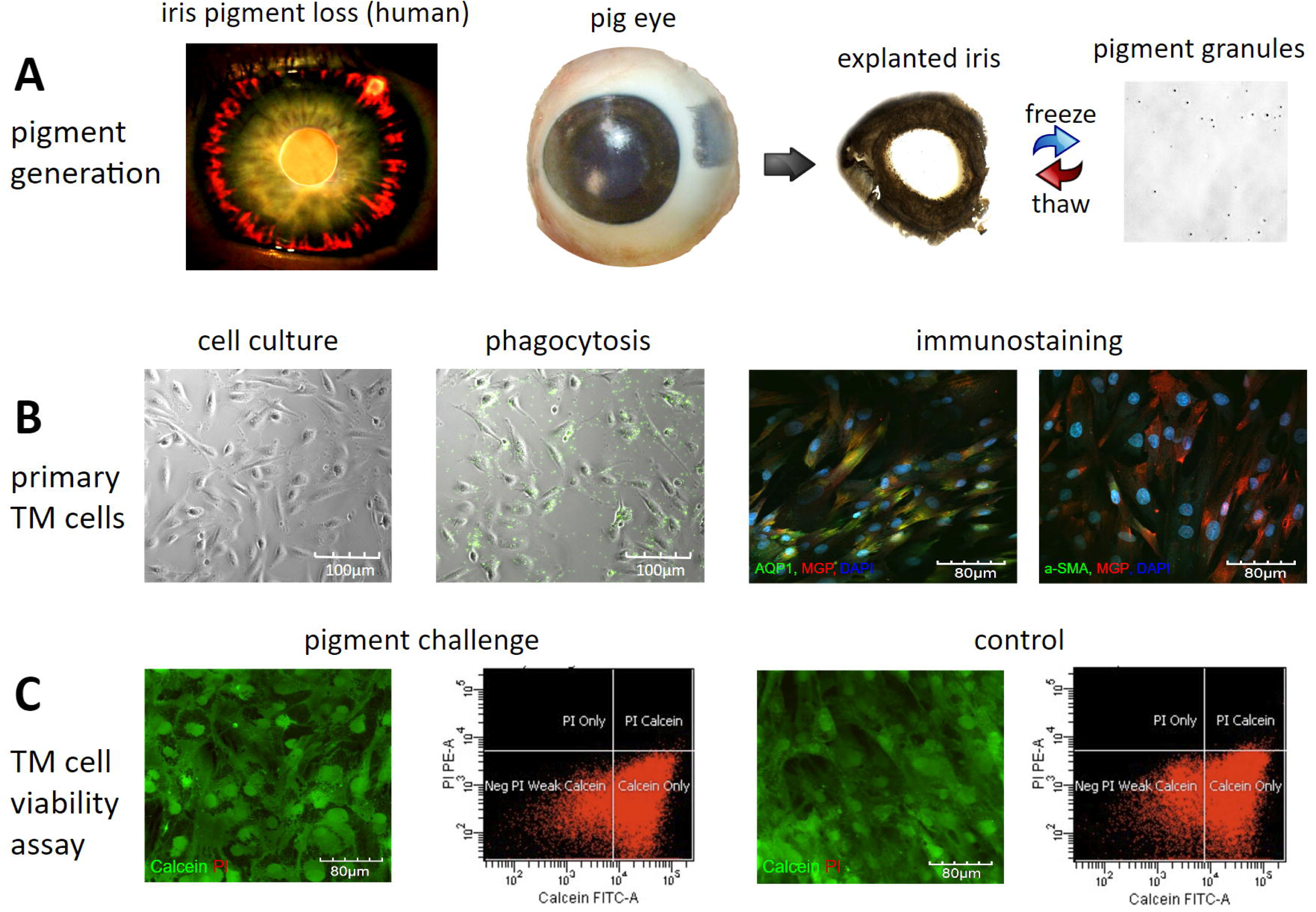
Pigment generation and in vitro exposure to pigment dispersion. In the human eye with pigment dispersion, pigment and stroma are lost in the mid-periphery of the iris (transillumination, A, left). Similar pigment granules can be generated by exposing an explanted pig iris to freeze-thaw cycles (A, middle and right). Granules had a size of 1.03±0.11 micron (A, right, single hemocytometer grid shown). Isolated primary trabecular meshwork (TM) cells from pig eyes (B, left to right) displayed the characteristic morphology, phagocytic activity (fluorescent microspheres) and immunostaining pattern with TM specific markers MGP, AQP1 and alpha-SMA (B, right). Exposure to pigment did not change the percentage of viable cells or propidium iodide-positive, dead or apoptotic cells (C).

Although the amount of pigment granules in the aqueous humor is correlated to IOP^8^, the observed amount^9^ is insufficient to cause a simple physical outflow obstruction as a primary mechanism. Different models of pigment dispersion exist: DBA/2J^10^ mice experience ocular hypertension as a result of synechial angle closure, iris atrophy and pigment dispersion^10^. In contrast, Col18a1(-/-) mice^11^ have a collagen XVIII/endostatin deficiency that leads to pigment dispersion through an unknown mechanism but lack ocular hypertension. Mouse eyes have a limited number of TM layers and are approximately 455 times smaller than human and porcine eyes^12^, making ex vivo cultures more challenging^13^. Monkeys can develop an IOP elevation from repeated intracameral pigment injections^14^ but bolus applications do not reflect the chronic pigment release in PG well and primate models make it costly to power a study with a sufficient number of animals. Bolus injections of pigment in normal rodent eyes would be difficult to accomplish due to the small anterior chamber volume of only a few microliters.

Our prior work and the study presented here, took advantage of the high tissue quality of pig eyes caused by a short time from enucleation to culture (two hours), the consistency within a litter, and an outflow tract anatomy that matches several features in humans^15–18^. Important differences are a thicker TM, Schlemm’s canal like segments instead of a mostly single lumen (angular aqueous plexus)^19^ and, in contrast to almost all other domestic animals and pets^20^, the virtual absence of naturally developing glaucoma or medically induced-ocular hypertension. We recently established gene transfer^17, 21^, modeled segmental aqueous outflow^16,22,23^ and created a microincisional angle surgery system^24–26^ in pig eye models.

Here, we hypothesized that ex vivo perfused pig eyes would experience a relative outflow failure from continuous exposure to pigment at a concentration far lower (10,000-fold) than used in previous bolus experiments^14^. The aim of this study was to develop a standardized and accessible PG model that allows the study of TM function and signal pathway changes to identify new treatment targets.

## Results

We developed a porcine eye model of PG that replicates the clinical features in human patients, consisting of a higher concentration of pigment granules in the aqueous humor^8^ and inside of TM cells^9,27,28^.

### Pigment granules do not affect the viability of primary TM cells

The freeze-thaw method produced pigment granules of 1.03±0.11 micron that were similar to the pigment in human pigmentary glaucoma. Stocks could be kept at a concentration of 4.3x10^9^ particles/ml without clumping **(Fig. 1A)**. Primary TM cells obtained from freshly prepared TM exhibited the characteristic flat, elongated trabecular shapes^29^ that became more spindle-tipped with increasing confluence and were positive to TM specific markers MGP^30–33^, AQP1^32, 34^ and alpha-SMA^35–37^ **(Fig. 1B)**.

The cytotoxicity of pigment granules was evaluated by flow cytometry and immunostaining using calcein acetoxymethyl (AM) and propidium iodide (PI) co-labelling. The non-fluorescent AM derivative of calcein is transported through the cellular membrane into live cells while PI cannot cross the membrane of live cells, making it useful to differentiate necrotic, apoptotic and healthy cells^38^. Viable TM cells can convert non-fluorescent calcein AM to green fluorescent calcein by intracellular esterase, but do not allow PI to enter or to bind nucleic acids^39^. Pigment granules at 1.67x10^7^ particles/ml did not increase the percentage of PI labelled apoptotic or dead cells (0.00±0.00% in the pigment group versus 0.27±0.07% in the normal control, P>0.05). Similarly, the percentage of calcein labelled viable cells was not decreased (84.90±3.87% in the pigment group compared to 84.57±3.00% in the normal control, P>0.05) **(Fig. 1C)**.

### Pigment elevates intraocular pressure

Eight left-right matched porcine anterior segment pairs were randomly assigned to the pigment or the control group, respectively. Similar baseline IOPs were established for 72 hours (11.80±0.78 mmHg versus 11.64±0.51 mmHg, *P*= 0.872), followed by 180 hours of pigment exposure at a concentration of 1.67 x 10^7^ particles/ml, or a sham treatment with normal tissue-culturing medium. The control group did not have any significant change of IOP throughout the experiment (all *P*>0.05, compared to the baseline). In contrast, pigment resulted in an IOP elevation that became significantly higher at 48 hours, peaked at 96 hours at 23.0±6.0 mmHg and persisted at an IOP that was 75% above the baseline of 11.5±0.8 mmHg (all *P*<0.05, compared to the group average baseline). Pigment dispersion eyes also had a significantly greater IOP fluctuation than controls (1.78±0.62 versus 0.83±0.22, *P*<0.001, mean±standard deviation), indicating different IOP responses of individual eyes to pigment treatment existed **(Fig. 2)**.

**Figure 2.**
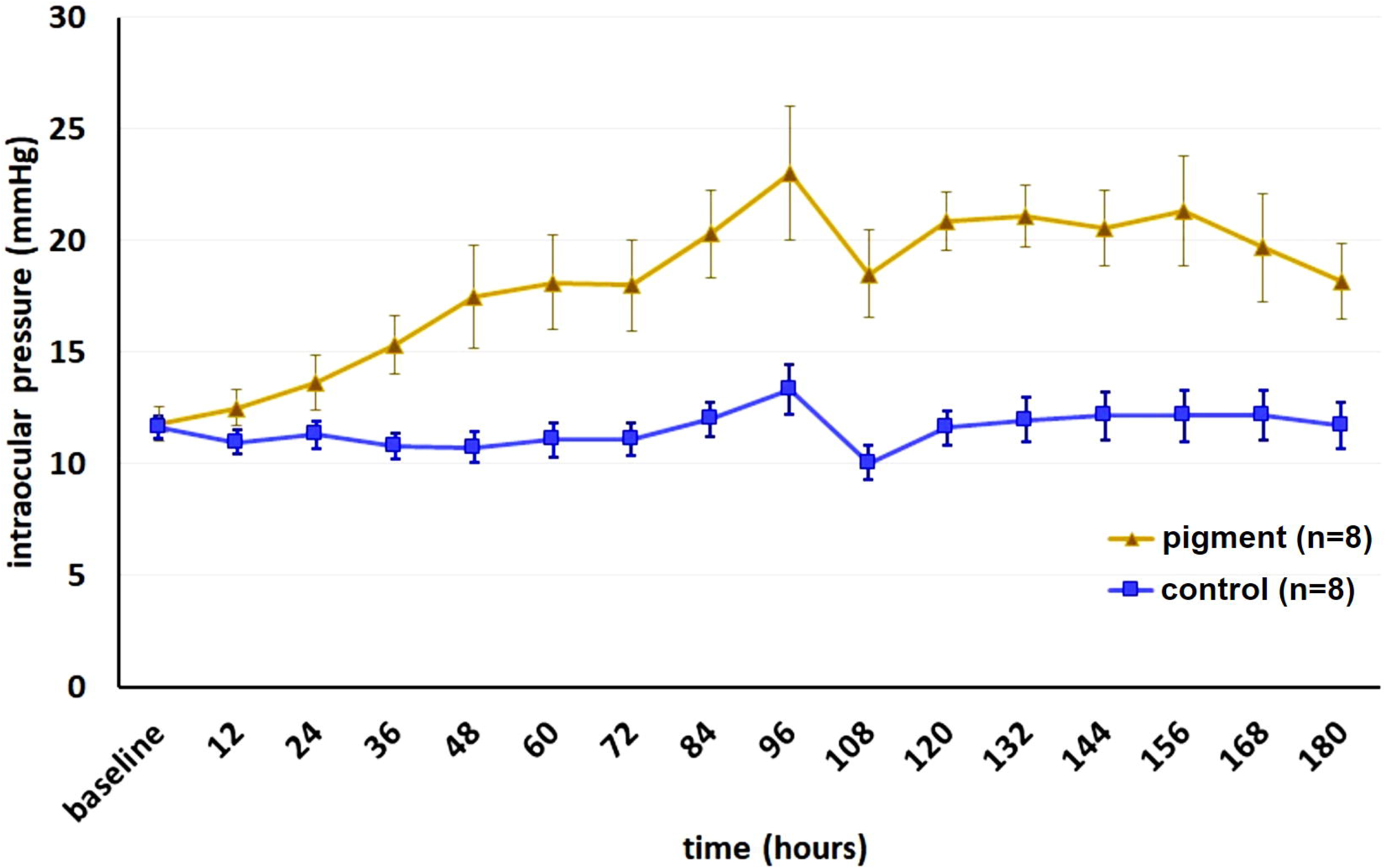
IOP elevation following pigment dispersion. IOP of perfused anterior segments started to increase after 48 hours (*P*=0.026) and persisted for the remainder of the study (all P<0.05). IOP fluctuation in the pigment group was significantly larger than in the paired controls (*P*<0.001). A medium change occurred after the 96 hour time point causing the parallel dip in IOP.

### Lysosome and phagosome activated by pigment

Anterior segments of the control eyes presented with a normal TM consisting of the uveal meshwork, corneoscleral meshwork, and juxtacanalicular region immediately adjacent to the inner wall of Schlemm’s canal like segments of the angular aqueous plexus **(Fig. 3A)**. Only occasional, scattered pigment was seen that was intracellular **(Fig. 3I, red arrowhead)**. In contrast, pigment dispersion eyes contained many TM cells with intracytoplasmic pigment granules, including enlarged cells that were protruding into elements of the downstream outflow tract **(Fig. 3B, C, F, G, J, K, red arrowheads)**. There was no evidence of a collapse of intertrabecular spaces or a physical pigment obstruction of the TM or collector channels **(Fig. 3B and C)**. Ultrastructurally, pigment induced the activation of lysosomes and phagosomes both ex vivo and in vitro **(Fig. 3K and L, blue arrowheads)**. Numerous pigment granules appeared ingested and at different stages of hydrolysis by secondary lysosomes **(Fig. 3K, and L)**. A swollen and distended endoplasmic reticulum could also frequently be seen in these eyes **(Fig. 3J, and L, yellow arrowheads)**.

**Figure 3.**
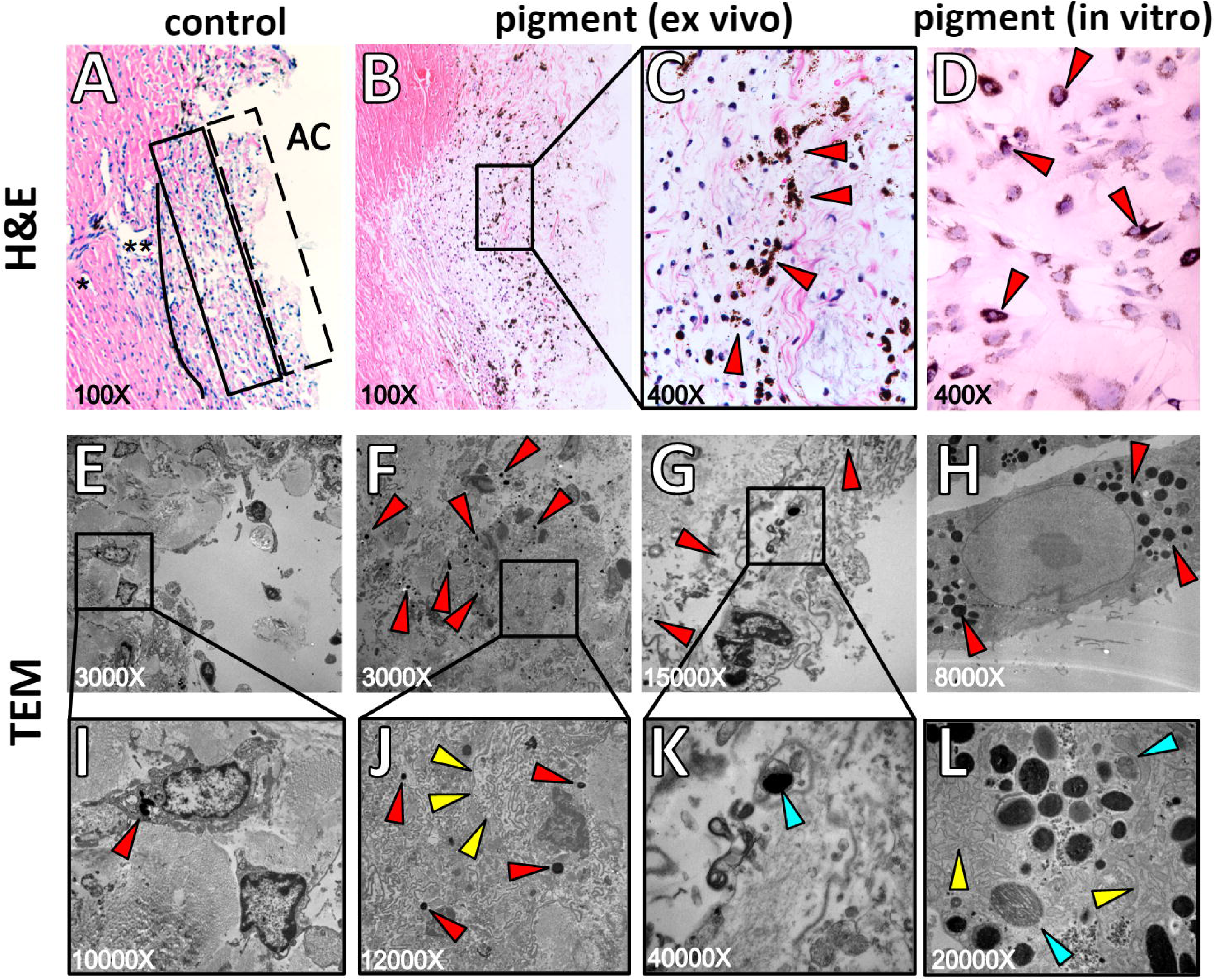
Ultrastructure and histology of the trabecular meshwork (TM). The TM consisted of three characteristic layers: the uveal meshwork (box with dashed line, Figure 3A), the corneoscleral meshwork (box with solid line, Figure 3A) and the juxtacanalicular meshwork (solid line, Figure 3A), adjacent to the inner wall of Schlemm’s canal (SC) (Figure 3A, black asterisks). The outer layers were phagocytically active. Pigment granules were located within cells and around the nucleus in the ex vivo model (Figure 3B and 3C, red arrows) and in vitro (Figure 3D, red arrows). Transmission electron microscopy showed occasional pigment in the normal TM (Figure 3E and 3I, red arrows), but a larger number in the inner TM layers (Figure 3F and Figure 3J, red arrows), the outer TM layers (Figure 3G and 3K, red arrows) and in primary TM cells after pigment treatment (Figure 3H). Pigment hydrolysis in different phagolysosome stages (Figure 3K and 3L, blue arrows) and endoplasmic reticulum (Figure 3J and 3L, yellow arrows) were also seen in vitro and ex vivo.

### Declining phagocytosis, and cytoskeleton disruption

The reorganization of the cytoskeleton^40^, reduced phagocytosis^41, 42^, features closely related to regulation of outflow^43^, were the most notable observations in the eyes exposed to pigment.

Primary TM cells normally have flat, elongated bodies with drawn out, spindle shaped tips before assuming a spindle shape with increasing confluence **(Fig. 4A1).** This was not significantly affected by the pigment treatment **(Fig. 4A2)**. We used phalloidin to label the F-actin cytoskeleton of TM cells. Normal F-actin fibers presented as well-organized, fine, feather-like microfilaments **(Fig. 4B1, white arrowheads)** while pigment caused the polymerization of F-actin fibers into long, thick bundles **(Fig. 4B2, red arrowheads)**. The quantitative analysis showed a significantly higher percentage of TM cells containing actin stress fibers in the pigment treated group than that of the control (32.64± 2.37% vs. 52.16± 1.69%, *P*<0.001) **(Fig. 4C)**.

**Figure 4.**
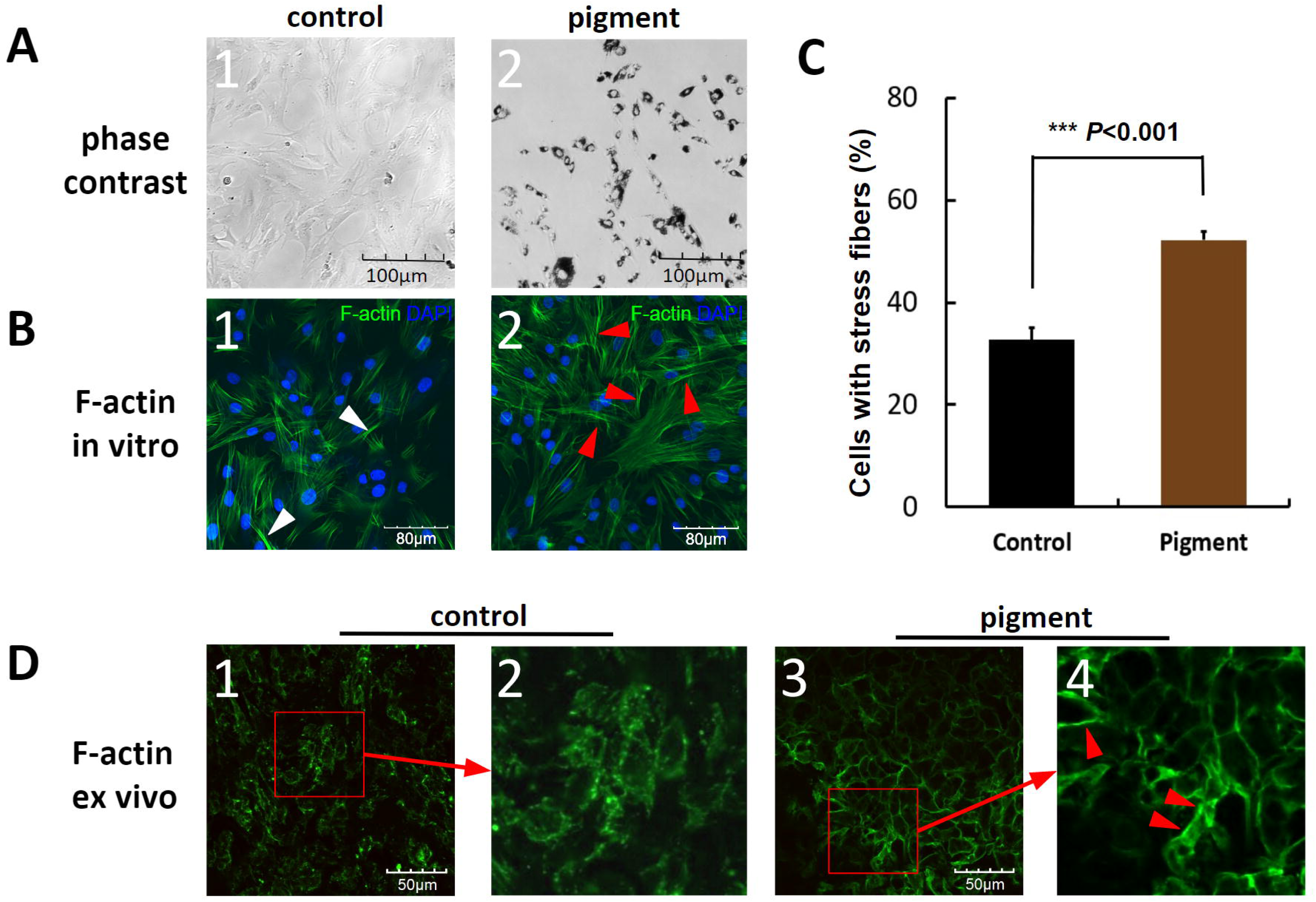
TM cytoskeletal changes induced by pigment. Primary TM cells (Figure 4A1) did not show morphological changes when exposed to pigment granules (Figure 4A2). Normal F-actin cytoskeleton (white arrowheads, Figure 4B1). Pigment induced actin stress fibers with long, thick bundles (red arrowheads, Figure 4B2). Actin stress fibers increased from 32.64± 2.37% in controls to 52.16± 1.69% in the pigment group (*P*<0.001). F-actin cytoskeleton of flat-mounted normal TM tissue with weak, segmental structures (Figure 4D1 and 4D2), in contrast to thick, continuous stress fibers in the pigment group (Figure 4D3 and 4D4).

Consistent with the in vitro findings, F-actin microfilaments in the normal TM flat-mounted tissue samples showed weak, spot-like or segmental stainings **(Fig. 4D1 and 2)**, contrasting the thick, bundle-like, continuous stress fibers in the pigment group **(Fig. 4D3 and 4)**.

In vitro phagocytosis was measured by flow cytometry. Normal primary TM cells readily phagocytosed carboxylate-modified green-yellow microspheres. In contrast, cells exposed to pigment had a 5.17 fold decreased uptake (**Fig. 5A**; controls: 48.7±2.17%, pigment dispersion eyes 9.4±4.2%, *P*<0.001). We also developed a semiquantitative method to measure the ex vivo TM phagocytosis. Carboxylate-modified microspheres were perfused into the anterior chambers to be phagocytosed by TM cells in situ.

**Figure 5.**
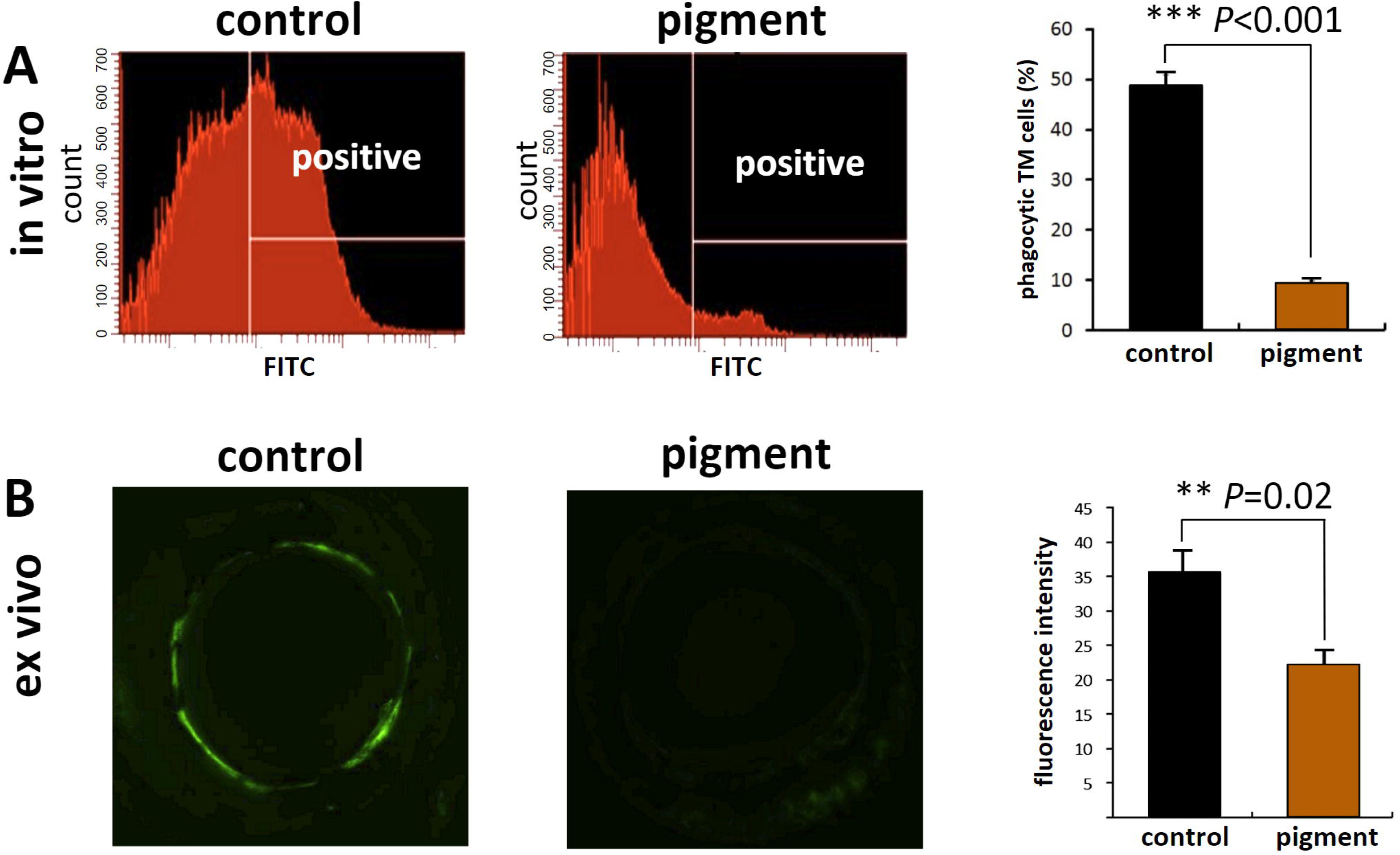
TM phagocytosis. In vitro TM phagocytosis was quantified by flow cytometry. Normal primary TM cells phagocytose fluorescent microspheres, but pigment dispersion reduces uptake 5.17-fold (48.73±2.17% versus 9.43±4.2%, P<0.001 (A)). Ex vivo TM phagocytosis was quantified in a similar fashion but by measuring fluorescence of inverted anterior segments instead (B, inverted culture dish with a direct view of the entire, fluorescent meshwork^17, 22^). Compared to the controls, fluorescence was significantly lower (3.4x10^7^±4.5x10^6^ versus 2.2x10^7^±2.1x10^6^, P= 0.020; **= P<0.01, ***= 0.001).

Fluorescent intensity could be observed with an epifluorescence equipped dissecting microscope only after uptake. The raw TM fluorescent intensity in the control group was 3.4×10^7^±4.5×10^6^, significantly higher than that of the pigment group 2.2×10^7^±2.1×10^6^ **(Fig. 5B**, *P*=0.020).

We also quantified cell-matrix adhesion as previously described^44^. Confluent TM monolayers that received pigment or vehicle treatment were subjected to trypsinization to measure cell-matrix adhesion. The numbers of TM cells per visual field showed no significant difference between the two groups before trypsinization (230.00±5.51 in the pigment group versus 244.33±6.39 in the controls, *P*=0.810). After trypsinization a higher number of TM cells in the pigment group started to contract, show shrinkage of the cell body, and detach. The remaining TM cells in the pigment group were significantly fewer than than in the control group at 2 minutes (173.33±10.81 versus 205.00±1.53), *P*=0.038) and 5 minutes (112.33±11.30 versus 158.67±6.94, *P*=0.010) **(Fig. 6)**. Cell migration was assessed by the numbers of cells which migrated into six-well plate from glass slides that were pre-populated with a TM monolayer. An average of 54,583±8,718 TM cells migrated onto the six-well plate after ten days of pigment treatment, in contrast to 33,000±5,500 cells in the control, but there was no statistical difference (*P* =0.231).

**Figure 6.**
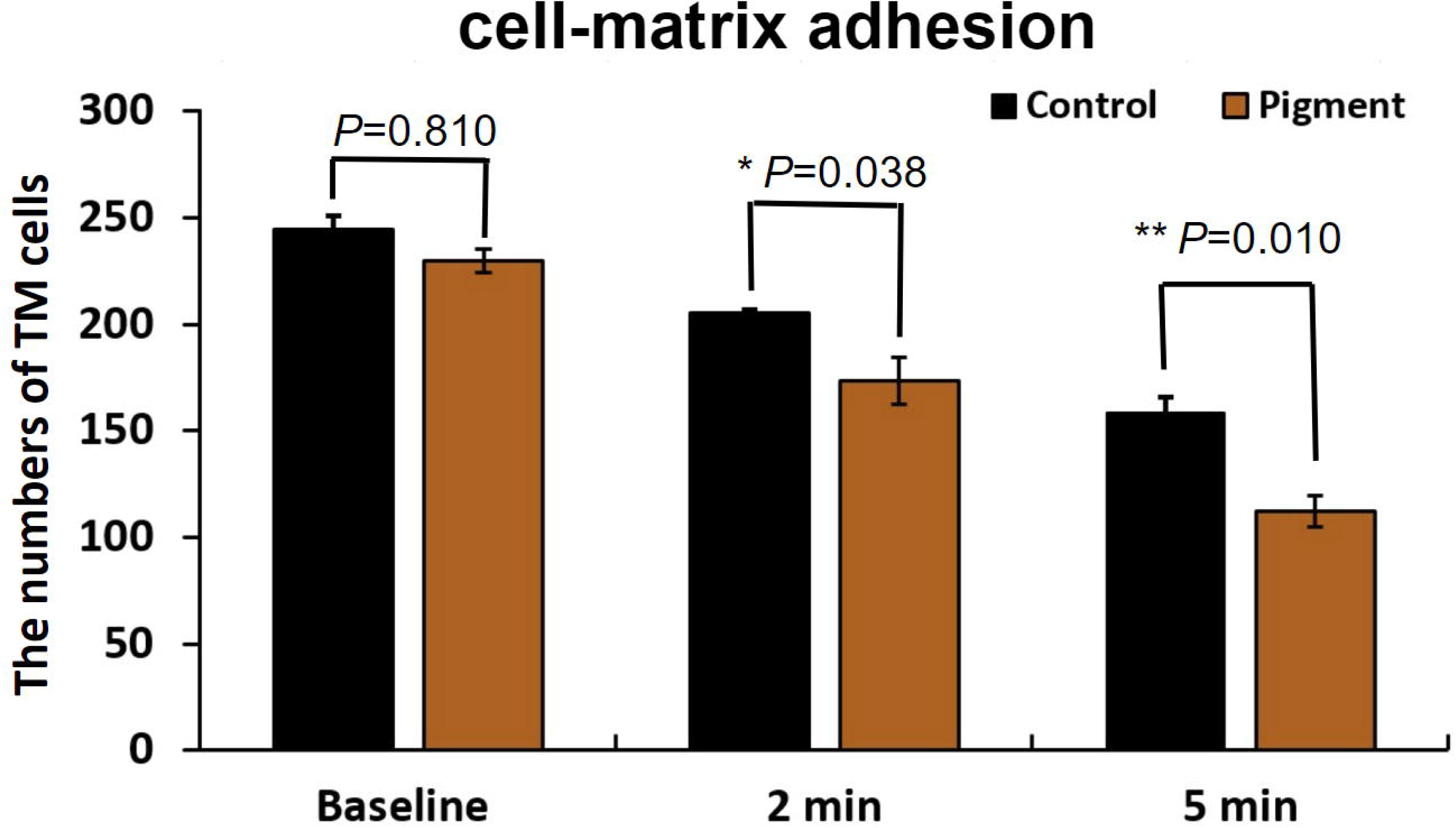
Cell matrix adhesion. In the pigment group, there were significantly fewer TM cells at 2 minutes (173.33±10.81 versus 205.00±1.53, P = 0.038) and 5 minutes (112.33±11.30 versus 158.67±6.94), *P* = 0.010), compared to the control (Figure 6, left).

### Pigment changes pathways of cellular movements, phagocytosis, and aqueous outflow

Three ex vivo TM samples from each group were submitted for analysis with a gene expression microarray. A total of 24,123 *porcine* genes were analyzed, of which 691 were upregulated **(red dots in volcano plot Fig. 7A and red lines in heatmap Fig. 7B)** and 332 were downregulated more than 1.5 fold **(green dots in volcano plot in Fig. 7A and green lines in heatmap Fig. 7B** and **Supplemental Table 1**, *P*<0.05). After excluding 239 porcine genes with unclear biological functions, 784 genes (**Supplemental Table 2**) were mapped in our pathway analysis to 16 distinct signaling pathways. They were related to (1) cellular movement (cell adhesion, diapedesis and migration), (2) endocytosis and phagocytosis, (3) aqueous outflow facility, (4) oxidative stress and endoplasmic reticulum stress, and (5) TM extracellular matrix remodelling **(Fig. 8)**. According to this pathway modelling, RhoA signaling, the central pathway that regulates the TM actin cytoskeleton is initialized by pigment exposure via a complex consisting of the insulin growth factor (IGF), the type 1 insulin-like growth factor receptor (IGF-IR) and the lysophosphatidic acid receptor (LPAR) in the cell membrane. This is different from TGFβ and its receptor-induced-RhoA activation in POAG^45^ and steroid glaucoma^46^. Besides the direct inhibition of tight junction formation by RhoA activation, upregulation of the rhophilin Rho GTPase binding protein (RHPN) would also promote the reorganization of the TM actin cytoskeleton. This negatively affects the tight junction protein 2/zonula occludens-associated nucleic acid binding protein complex (TJP2/ZONAB), and inhibits TM tight junction and clathrin, caveolae or Fc? receptor-mediated endocytosis and phagocytosis. The pathway analysis also indicated that activation of RhoA signaling enhances myosin/MYBPH-mediated actin polarization, stress fiber formation and TM contractility which affects TM motility.

**Figure 7:**
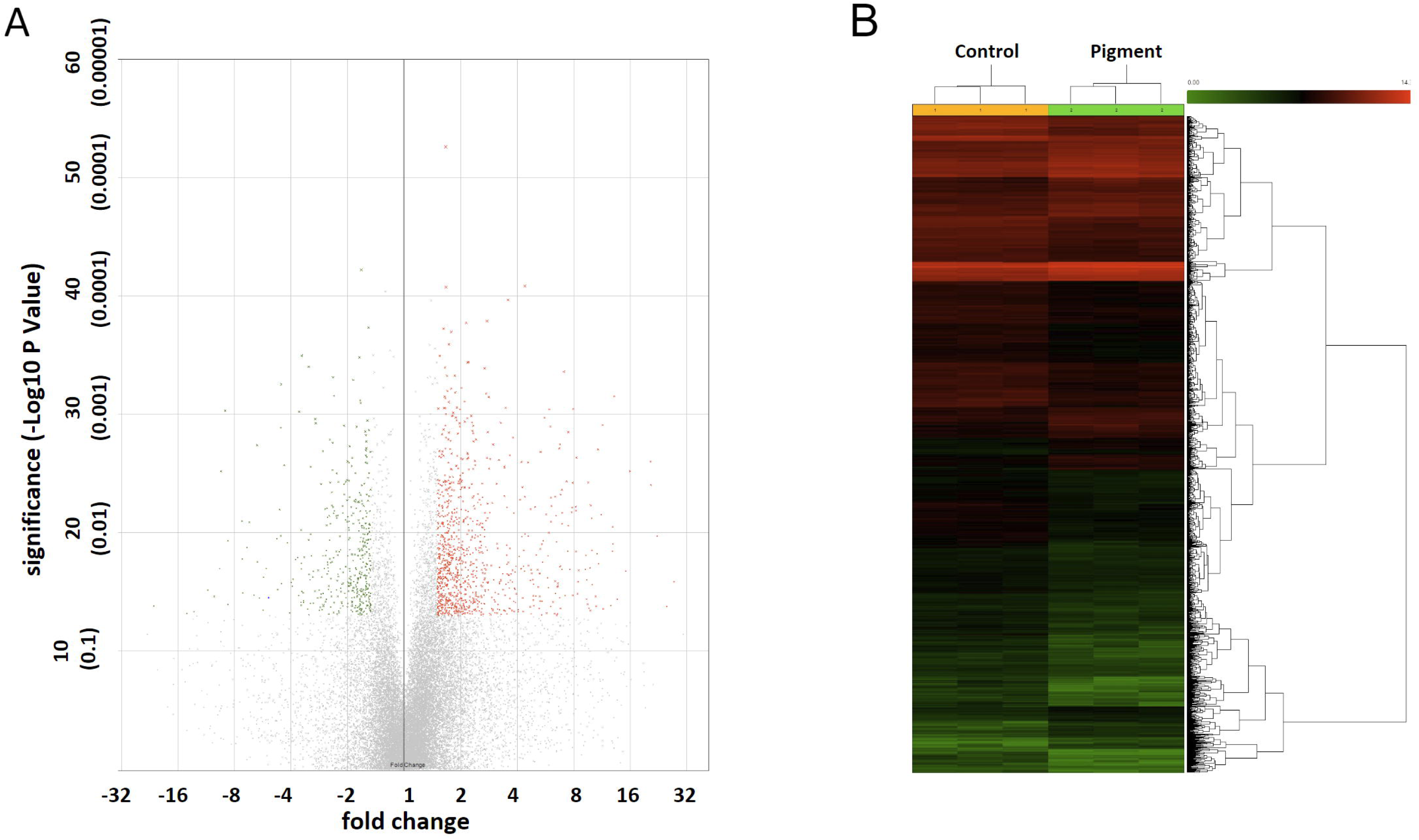
Differential gene expression by pigment treatment. Three TM samples from eyes that had a confirmed IOP elevation phenotype were compared to controls using the Affymetrix Gene 3’ IVT Microarray. A total of 24,123 porcine genes were hybridized, of which 691 were upregulated (red dots in volcano plot and red lines in heatmap) and 332 were downregulated (green dots in volcano plot and green lines in heatmap) by more than 1.5 fold with p value ≤0.05.

**Figure 8.**
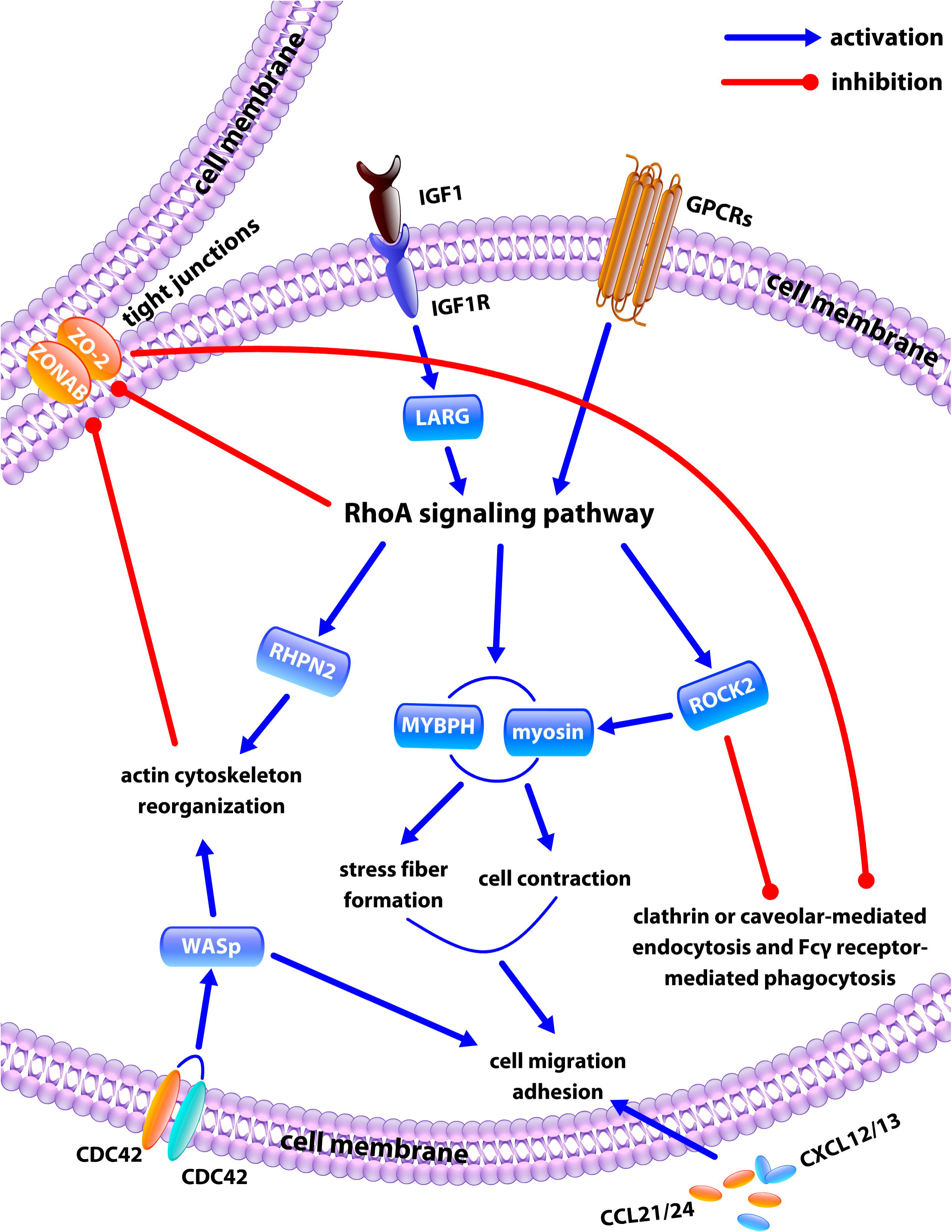
Pathway analysis of trabecular meshwork exposed to pigment. RhoA signal is initialized by a complex consisting of the insulin growth factor (IGF), the type 1 insulin-like growth factor receptor (IGF-IR) and the lysophosphatidic acid receptor (LPAR).

Additionally, change of TM motility could also result from the upregulation of a set of chemokine ligands (CCL21/CCL24 and CXCL12/CXCL13) in the cell membrane and Wiskott-Aldrich syndrome protein (WASp) in the cytoplasm. Key genes and signal pathways are summarized in **Table 1** while their upstream regulators are listed in **Supplemental Table 3**. Genes involved with cellular movement were significantly upregulated **(Supplementary Figure 1)**.

**Table 1:** Expression changes of key genes and their pathways after exposure to pigment as indicated by pathway analysis.

| Gene symbol | Entrez gene name | Fold change | P-value (ANOVA) | Signal pathways | Biological functions |
| --- | --- | --- | --- | --- | --- |
| **LPAR3** | lysophosphatidic acid receptor 3 | 6.02 | 0.042 | eNOS signal pathway, VEGF signal pathway, RhoA signal pathway and Gα12-13 signal pathway | Outflow regulation |
| **PGF** | placental growth factor | 2.49 | 0.00164 | Ephrin receptor signal pathway, VEGF signal pathway, eNOS signal pathway, clathrin-mediated endocytosis signal pathway, IL-8 signal pathway, and hepatic fibrosis signal pathway | Outflow regulation; extracellular matrix remodeling; cell adhesion and migration |
| **PIK3R2** | phosphoinositide-3-kinase regulatory subunit 2 | 2.13 | 0.0389 | IL-8 signal pathway, VEGF signal pathway, eNOS signal pathway, NRF2-mediated oxidative stress response signal pathway, Gα12-13 signal pathway, clathrin-mediated endocytosis signal pathway, Fcγ receptor-mediated phagocytosis signal pathway, JAK-STAT signal pathway | Outflow regulation; cell migration; cell phagocytosis; oxidative stress and endoplasmic reticulum stress |
| **ALB** | albumin | 2.11 | 0.0299 | Caveolar-mediated endocytosis signal pathway, and clathrin-mediated endocytosis signal pathway | Cell endocytosis |
| **WAS** | Wiskott-Aldrich syndrome | 2.1 | 0.0439 | Ephrin receptor signal pathway, Fcγ receptor-mediated phagocytosis signal pathway | Cell adhesion and migration |
| **CDH5** | cadherin 5 | 2.05 | 0.0119 | Agranulocyte adhesion and diapedesis signal pathway, and Gα12-13 signal pathway | Cell adhesion; outflow regulation |
| **KDR** | kinase insert domain receptor | 2.02 | 0.000598 | eNOS signal pathway, and VEGF signal pathway | Outflow regulation |
| **FLT1** | fms related tyrosine kinase 1 | 1.98 | 0.0319 | eNOS signal pathway, and VEGF signal pathway | Outflow regulation |
| **ITGB8** | integrin subunit beta 8 | 1.96 | 0.0121 | Caveolar-mediated endocytosis signal pathway, and clathrin-mediated endocytosis signal pathway | Cell endocytosis |
| **CXCL12** | C-X-C motif chemokine ligand 12 | 1.9 | 0.0264 | Agranulocyte adhesion and diapedesis signal pathway, and ephrin receptor signal pathway | Cell adhesion and migration |
| **IGF1-R** | insulin like growth factor 1 receptor | 1.8 | 0.0165 | PETN signal pathway, RhoA signal pathway, hepatic fibrosis signal pathway | Extracellular matrix remodeling; cell adhesion; outflow regulation |
| **PIK3C2B** | phosphatidylinositol-4-phosphate 3-kinase catalytic subunit type 2 beta | 1.64 | 0.0118 | IL-8 signal pathway, VEGF signal pathway, eNOS signal pathway, NRF2-mediated oxidative stress response signal pathway, Gα12-13 signal pathway, clathrin-mediated endocytosis signal pathway, JAK-STAT signal pathway | Outflow regulation; cell migration; cell endocytosis; oxidative stress and endoplasmic reticulum stress |
| **LPAR6** | lysophosphatidic acid receptor 6 | 1.56 | 0.0456 | eNOS signal pathway, VEGF signal pathway, RhoA signal pathway, and Gα12-13 signal pathway | Outflow regulation |
| **IGF-1** | insulin like growth factor 1 | 1.54 | 0.0452 | Clathrin-mediated endocytosis signal pathway, RhoA signal pathway, hepatic fibrosis signal pathway | Extracellular matrix remodeling; cell endocytosis; outflow regulation |
| **FYN** | FYN proto-oncogene, Src family tyrosine kinase | -1.51 | 0.0454 | Ephrin receptor signal pathway, Fcγ receptor-mediated phagocytosis signal pathway, caveolar-mediated endocytosis signal pathway | Cell adhesion and migration; cell phagocytosis; cell endocytosis |
| **ROCK2** | Rho associated coiled-coil containing protein kinase 2 | -1.56 | 0.0116 | Ephrin receptor signal pathway, RhoA signal pathway, VEGF signal pathway, Gα12-13 signal pathway, IL-8 signal pathway | Cell migration; cell adhesion; outflow regulation |
| **CDH3** | cadherin 3 | -1.56 | 0.0114 | Gα12-13 signal pathway | Outflow regulation |
| **PKC-Z** | protein kinase C zeta | -1.75 | 0.0488 | IL-8 signal pathway, PETN signal pathway, eNOS signal pathway, NRF2 mediated oxidative stress response signal pathway, Fcγ receptor-mediated phagocytosis signal pathway | Cell migration; cell adhesion; cell phagocytosis; oxidative stress and endoplasmic reticulum stress; outflow regulation |
| **ITGA6** | integrin subunit alpha 6 | -1.77 | 0.0458 | Agranulocyte adhesion and diapedesis signal pathway, and caveolar-mediated endocytosis signal pathway | Cell adhesion; cell endocytosis |
| **ITGA2** | integrin subunit alpha 2 | -1.8 | 0.00624 | Ephrin receptor signal pathway, caveolar-mediated endocytosis signal pathway | Cell adhesion and migration |
| **VEGFA** | vascular endothelial growth factor A | -2.45 | 0.0244 | VEGF signal pathway, eNOS signal pathway, clathrin-mediated endocytosis signal pathway, ephrin receptor signal pathway, IL-8 signal | Extracellular matrix remodeling; cell adhesion and migration; cell |

|  |  |  |  | pathway, hepatic fibrosis signal pathway | endocytosis; outflow regulation |
| --- | --- | --- | --- | --- | --- |
| **DNAJC3** | DnaJ heat shock protein family (Hsp40) member C3 | -2.56 | 0.00741 | Unfolded protein response signal pathway, NRF2-mediated oxidative stress response signal pathway | Oxidative stress and endoplasmic reticulum stress; oxidative stress and endoplasmic reticulum stress |
| **DNAJB9** | DnaJ heat shock protein family (Hsp40) member B9 | -3.12 | 0.0157 | Unfolded protein response signal pathway, NRF2-mediated oxidative stress response signal pathway | Oxidative stress and endoplasmic reticulum stress; oxidative stress and endoplasmic reticulum stress |
| **CLDN2** | claudin 2 | -11.08 | 0.0305 | Agranulocyte adhesion and diapedesis signal pathway | Cell adhesion; |

## Discussion

In this study, we developed a porcine ex vivo model of PG, a common and often risky type of secondary glaucoma in humans, to study the TM function and signal pathway changes. To the best of our knowledge, this is the first ex vivo system to do so in any species. In analogy to a freeze-thaw decellularization technique we recently developed^47^, we used a freeze-thaw technique to produce pigment particles from the iris while preserving epitopes and avoiding chemicals. In PG, iridozonular contact and pupillary movement^48^ cause disruption and loss of cells within the iris stroma and the pigmented iris epithelium^49^. The debris also contains melanin containing melanosomes^50, 51^. Our pigment granules had a size which matches those seen in human PG well^52^. They induced a hypertensive IOP phenotype without a physical obstruction to the intertrabecular spaces, similar to that in human PG patients^47^. As discussed in the following, our results suggest that ocular hypertension in PG is the result of actin stress fibers, reduced phagocytosis and cell-matrix adhesion.

We isolated primary porcine TM cells for in vitro experiments. These TM cells displayed the typical TM markers^53^ seen in human TM cells and readily phagocytosed particulate matter. When their viability was not affected when exposed to pigment we proceeded to recreating pigment dispersion in our ex vivo perfusion model.

IOP started to elevate in eyes with continuous pigment dispersion at 48 hours and remained at a similar level thereafter. These eyes also displayed a larger range of IOPs compared to the normal controls, indicating a range of individual IOP responses, an observation that correlates to the IOP in pigment dispersion syndrome and PG in human patients^28^. The IOP elevation of 10 mmHg above baseline is comparable to a typical clinical setting^3, 4^.

The ultrastructure and histological findings matched studies of human eyes^9,27,54,55^. Most importantly, there was no physical obstruction to outflow but rather a relatively modest deposition of pigment within the TM and its cells. In the anterior segments with pigment diffusion, TM cells were busy with phagocytosis and breakdown of ingested pigment granules. The TM cytoskeleton displayed stress fibers with polymerization of F-actin microfilaments. Different from the formation of cross-linked actin networks in other forms of glaucoma^56^, for instance, steroid induced glaucoma^57^ and POAG^58^, the stress fibers in this model were long, thick, bundle-like microfilaments. Human TM cells could form cross- linked actin networks (CLANs) within seven days in steroid-induced ocular hypertension^56, 57^. An acute disruption of F-actin was reported in latex microspheres-induced phagocytic challenge of bovine TM cells^59^. TM cytoskeletal changes that directly affect the TM stiffness and outflow facility^60^ have recently been targeted with newer glaucoma medications in the form of ROCK inhibitors^61^ and NO donors^62^ that relax the TM. In contrast, dexamethasone^63^, TGF-β2^64^ and senescence^65^ result in an increased extracellular matrix stiffness and decreased outflow.

The TM does not only regulate outflow but also prevents debris from entering the outflow system but has the ability to phagocytose it and present it to the immune system^66, 67^. Genes for cell motility were significantly upregulated possibly reflecting migration of TM cells into SC as has been reported in human PG^9, 27^. Conversely, we observed a decline of cell-matrix adhesions and altered cytoskeleton as reported in another study^68^. It is possible that a limited amount of pigment may cause no harm but when a threshold is exceeded, the phagocytic capacity is overwhelmed as evidenced by reduced uptake of fluorescent spheres. TM cells may then migrate off the extracellular matrix of the TM and potentially occlude the intertrabecular space or the downstream outflow tract. This may explain why the pathology of TM in pigmentary glaucoma is so different from that in pigment dispersion syndrome^28^. Gottanka et al^9^ found that the Schlemm’s canal in human PG specimens was partially (25%- 65%) obstructed by loosely arranged, distended juxtacanalicular TM cells.

The gene expression changes in this model indicated a central role for RhoA and its activators IGF/IGF1R/LPAR in the molecular pathogenesis of PG with altered actin cytoskeleton and cell motility^69^. Activation of the RhoA pathway is seen in primary open angle^45^ and in steroid induced glaucoma glaucoma as well^46^. In contrast to TGFβ mediated RhoA activation in POAG^45^ and steroid-induced glaucoma^46^, this process seems to be initiated by IGF1. IGF1 has been implicated in ocular neovascularization^70^ and it is tempting to speculate about a role in the uveitis-hyphema-glaucoma (UGH) syndrome^71^. In UGH, intraocular lens implants cause chronic pigment release, iris atrophy, and hyphema.

TM tight junctions are formed by the interaction between actin fibers and tight junction proteins. Junctional tightness and distribution influence aqueous outflow facility and cell adhesion^72^. The downregulation of CDLN2 may be related to increased cell migration^73^. Activation of RhoA caused a disruption of TM tight junctions by modulating actin stress fibers^74, 75^. The downregulation of CLDN2 and ZO-2 may be related to increased cell migration and can also seen after steroid exposure^75^. Normal tight junctions also contribute to maintaining cell polarity, thereby allowing specialized surface functions such as receptor-mediated phagocytosis and endocytosis. Consistent with this downregulation, we found that clathrin or the caveolar-mediated endocytosis pathways and the Fcγ receptor-mediated phagocytosis pathway were all significantly downregulated by pigment exposure.

Compared to small rodent models, the advantages of the pig eye model presented here that eyes can be cultured ex vivo^17^, are readily available from local abattoirs^16,17,22,23,26,47^, TM can be observed directly through the bottom of the culture dish^17^, agents (vectors, pigment, drugs) can be applied continuously, techniques to measure regionally discrete outflow exist^16, 23^, ample trabecular meshwork material from a single animal (helpful in gene expression microarrays) and that a potential immune response to injection of material from other animals^14^ is eliminated ex vivo. Regardless, this model has several limitations. Compared to human PG, the IOP elevation was relatively acute and took two days from pigment exposure to ocular hypertension. In contrast, development of high IOP in human patients can take years^76, 77^. Debris clearing by macrophages, in addition to TM cell phagocytosis, is a more effective process in vivo than ex vivo^66, 67^. Although the accumulation of pigment granules are visually striking both microscopically and macroscopically, this does not mean that melanin itself is the principal cause of PG. To determine what cellular fragments, organelles or proteins contribute the most, perfusion experiments with different phases from a fractionation centrifugation would have to be performed.

In summary, we developed a PG model that manifested a hypertensive IOP phenotype as well as histological and ultrastructural characteristics similar to that of human PG. Pathway analysis revealed that activation of the RhoA pathway played a central role in the TM actin cytoskeleton, tight junction formation, phagocytosis, and cell motility, each of which might provide a target for PG treatment.

## Methods

### Generation of pigment

This study was designated by the Institutional Review Board of the University of Pittsburgh exempt status under section 45 CFR 46.101(b)(1) of the Code of Federal Regulations. Within 2 hours of sacrifice, ten fresh pig eyes obtained from a local abattoir (Thoma Meat Market, Saxonburg, PA) were decontaminated for 2 minutes with 5% povidone-iodine ophthalmic solution (betadine 5%; Alcon, Fort Worth,TX,USA) and hemisected. After removal of the posterior segment, lens, and ciliary body, the anterior segments were saved in plain DMEM for mounting. Irises were collected to produce pigment granules with two freeze-thaw cycles. Briefly, ten irises were placed in 15 ml PBS and frozen at -80°C for two hours, then thawed at room temperature. After two cycles of freeze-thaw, the tissue became fragile, and shed pigment when pipetted up and down 100 times with a 3ml Pasteur pipette. Pigment granules were filtered through a 70 µm cell strainer (cat#431751, Corning Incorporated, Durham, NC). After spinning at 3000 rpm for 15 minutes, the supernatant was discarded, and pigment granules were resuspended in 15 ml PBS and centrifuged again. Finally, the pellet was resuspended in 4 ml PBS and stored at 4°C as a stock suspension.

The concentration of pigment granules was measured by hemocytometer counts (cat#1490, Hausser Scientific, Horsham, PA)^14^. The stock suspension was diluted 1000-fold and visualized by a phase contrast microscope at 600× magnification (Eclipse TE200-E, Nikon Instruments Inc., Melville, NY). Particle size was calculated as the average of 100 particles.

### TM primary culture

A glaucoma surgeon experienced in microincisional glaucoma surgery^24–26,78–82^ carefully excised the TM from freshly enucleated pig eyes under an ophthalmic operating microscope (Stativ S4, Carl Zeiss, Oberkochen, Germany). It was sectioned into 0.5 by 0.5 mm pieces, and maintained in TM medium (OptiMEM (31985-070, Gibco, Life technologies, Grand Island, NY) supplemented with 5% FBS and 1% antibiotics (15240062, Thermo Fisher Scientific, Waltham, MA) in a 37°C humidified CO_2_ incubator. Primary TM cells migrated out of the tissues and formed clones on day 5. After 100% confluence, the cells were trypsinized for 5 minutes, centrifuged at 1000 rpm for 3 minutes and replated at a 1:3 ratio. Only the first four passage of the cells were used to reduce the chance of differentiation^83^.

Primary TM cells were authenticated both by immunostaining with TM specific antibodies and phagocytosis testing. Briefly, the cells were fixed in 4% PFA for 1 hour, washed with PBS three times, and incubated with the following primary antibodies at 4°C overnight: goat polyclonal matrix gla protein (MGP) antibody (1:100 dilution in PBS, sc-32820, Santa Cruz, Dallas, Texas), rabbit polyclonal anti alpha smooth muscle actin (alpha-SMA) (1:100, ab5694, Abcam, Cambridge, MA) and aquaporin 1 (AQP1) antibodies (1:100, Sc-20810, Santa Cruz, Dallas, Texas). After three rinses of PBS, donkey-anti-goat Alexa Fluor 647 (1: 1000, ab150131, Abcam, Cambridge, MA) and goat anti-rabbit IgG secondary antibodies were added for 45 minutes at room temperature. Cell nuclei were counterstained with DAPI (D1306, Thermo Fisher Scientific, Waltham, MA). Pictures were taken by an upright laser scanning confocal microscope at 400 ×magnification (BX61, Olympus, Tokyo, Japan).

TM phagocytic activity was visualized and quantified. Briefly, the cells at 80% confluency were incubated for one hour with carboxylate-modified yellow-green fluorescent microspheres (0.5 micron diameter, cat# F8813, Thermo Fisher, Waltham, MA) at a final concentration of 5x10^8^ microspheres/ml. This was followed by three rinses with prewarmed PBS. To visualize TM phagocytosis, pictures were taken with a phase contrast fluorescence microscope at 200× magnification. To quantify the TM phagocytic activity, the cells were trypsinized, and resuspended in 500 µl PBS for flow cytometry. The percentage of TM cells that had ingested fluorescent microspheres was determined.

### Cell viability assay

Primary TM cells were plated onto a six-well-plate at 1x10^5^ cells per well and maintained in TM medium containing 1.67x 10^7^ pigment granules per ml. TM medium without pigment served as a control. The medium was changed every three days for up to 10 days. Cells were stained with calcein-AM (0.3 µM, C1430, Thermo Fisher, Waltham, MA) and PI (1 µg/ml, P1304MP, Thermo Fisher, Waltham, MA) for 30 minutes, followed by trypsinization and resuspension into 500 µl PBS for flow cytometry. Viable TM cells have intracellular esterase activity which can convert non-fluorescent calcein AM to green fluorescent calcein, but do not allow red fluorescent PI to pass through the intact cell membrane and bind to the cell nucleus^39^. Accordingly, calcein-labeled TM cells were counted as viable while PI-stained cells assumed to be dead or apoptotic.

### Pigmentary glaucoma model

For the ex vivo perfusion model, fresh pig anterior segments were mounted and perfused with DMEM supplemented with 1% FBS and 1% antibiotics at a rate of 3 µl/min as described previously^17, 47^. IOP was measured intracamerally by a pressure transducer (SP844; MEMSCAP, Skoppum, Norway) and recorded by a software system(LabChart; ADInstruments, Colorado Springs, CO, USA)^17, 47^. The transducers were calibrated with a transducer tester (Veri-Cal; Utah Medical Products, Midvale, UT, USA). Baseline IOPs were obtained after 72 hours. Pigment granules diluted with perfusion medium to a concentration of 1.67x10^7^ particles/ml were perfused for 180 hours. Normal medium without pigment granules served as a control. IOPs were recorded at two-minute intervals.

For in vitro studies, primary TM cells were treated with the pigment containing medium and while the control group was sham treated. Briefly, 3x10^5^ primary TM cells were plated onto a 60-mm dish and maintained with OptiMEM supplemented with 5% FBS and 1% antibiotics for 24 hours. Pigment granules were then added to a final concentration of 1.67x10^7^ particles/ml. The medium was changed every three days. Normal TM medium without pigment served as a control.

### Histology and transmission electron microscopy (TEM)

Anterior segments from perfusion cultures were fixed with 4% paraformaldehyde for 24 hours, paraffin embedded, cut into 5µm sections, and stained with hematoxylin and eosin for histology^17^. The ultrastructure of ex vivo TM tissue and in vitro cell monolayers were evaluated by transmission electron microscopy (TEM). Preparation of TEM samples followed a previous protocol with minor modification^84^. The samples were prefixed with 2.5% glutaraldehyde in 0.05 M cacodylate buffer for 24 hours, washed with PBS three times and then postfixed with 1% osmium tetroxide solution overnight. After three rinses of PBS, samples were dehydrated with an increasing ethanol series (30%, 50%, 70%, 90% and 100% ethanol, 45 minutes each), followed by embedding in epon resin (Energy Beam Sciences, East Granby, CT). The epon was exchanged completely every hour for three hours and blocks were cured for two days at 60°C. Sections of 300 nm were obtained with a Reichert-Jung Ultracut 701701 Ultra Microtome and stained with 0.5% Toluidine Blue O Solution (S25613, Thermo Fisher Scientific, Waltham, MA). Ultrathin sections of 65 nm were obtained and placed on grids. After staining with uranyl acetate and lead citrate, pictures were taken under an 80 kV Jeol transmission electron microscope (Peabody, MA) at various magnifications.

### Phagocytic activity

The primary TM cells exposed to pigment granules or sham were incubated with fluorescent microspheres for 1 hour, washed with PBS three times, trypsinized, resuspended and then subjected for flow cytometry. The phagocytosis was quantified by measuring TM fluorescent intensity after perfusion with carboxylate-modified yellow-green fluorescent microspheres for 24 hours. Briefly, 0.5 µm carboxylate-modified yellow-green fluorescent microspheres at 5x10^8^/ml were added to the perfusion system for 24 hours. Anterior chambers were washed three times with PBS. TM cells that had phagocytosed microspheres showed bright green fluorescence under a dissecting fluorescence microscope (SZX16, Olympus, Tokyo, Japan). Images were acquired at a 680 x 510 pixel resolution and a 1/17 second exposure. The raw TM fluorescent intensity was measured by ImageJ as previously described (Version 1.50i, NIH)^26, 85^.

### TM cytoskeleton

F-actin was used to assess the cytoskeletal changes of TM. Ex vivo and in vitro TM samples were fixed by 4% PFA for 1 hour and washed with PBS three times. The samples were incubated with Alexa Fluor 488 phalloidin (1:40 dilution, A12379, Thermo Fisher, Waltham, MA) for 30 minutes and counterstained with DAPI. Images were acquired with an upright laser scanning confocal microscope at 600-fold magnification (BX61, Olympus, Tokyo, Japan).

### TM motility

The cell-matrix adhesion was evaluated using a previously described protocol^44^. Confluent TM monolayers treated with pigment granules or sham were washed with PBS and dissociated with 0.25% trypsin. The changes in cell morphology and adhesion were monitored by a phase-contrast microscope at different trypsinization time intervals (0, 2, and 5 minutes). The ratio of the number of attached TM cells to total TM cells represented the cell-matrix adhesion.

Cell migration: primary TM cells were plated onto 18 mm^2^ coverslips (2855-18, Corning Incorporated, Durham, NC). After 100% confluency, these cover slides were transferred to six-well plates and maintained in TM medium with or without pigment granules (1.67 x 10^7^ particles/ml). The medium was replaced every three days. Cover glasses were removed after ten days. Cells that migrated to the well from the cover glass were trypsinized and counted.

### Gene expression microarray and pathway analysis

Anterior segments (n=3 each) from the pigment treated and normal control groups were dissected after the IOP phenotypes were obtained. Cells from the TM were lysed with trizol (15596026, Invitrogen, Thermo Fisher, Waltham, MA) and sent to the Genomic Core Facility of the University of Pittsburgh. Amplification and hybridization were performed using the Affymetrix Porcine 3’IVT Array (900624, Affymetrix, Santa Clara, CA) which contains 23,937 probe sets to interrogate 23,256 transcripts in pig representing 20,201 *Sus scrofa* genes. The Affymetrix CEL files were extracted, normalized, and statistically analyzed by ANOVA using the Transcriptome Analysis Console (TAC, version 3.1, Affymetrix, Santa Clara, CA). The differential gene expression profiles were characterized by a volcano plot and heatmap. The default filter criteria were fold Change (linear) less than -1.5 or fold Change (linear) above 1.5, and ANOVA p-value smaller than 0.05. Genes that matched these criteria were selected for bioinformatic pathway analysis using Ingenuity Pathway Analysis (Spring Release 2017, Qiagen, Hilden, Germany).

### Statistics

Data was presented as the mean ± standard error to capture the uncertainty around the estimate of the mean measurement and to allow computation of the confidence interval. Differential gene expression was analyzed using TAC (Version 3.1, Affymetrix, Santa Clara, CA). Other quantitative data was processed by one-way ANOVA using PASW 18.0 (SPSS Inc., Chicago, IL, USA). A difference was considered to be statistically significant if p<0.05.

### Data availability statement

All data generated or analysed in this study are included in this published article and its Supplementary Information files.

## Acknowledgements

This study was supported by NEI K08-EY022737 (Nils A Loewen), the Initiative to Cure Glaucoma of the Eye and Ear Foundation of Pittsburgh (Nils A Loewen), the Wiegand Fellowship of the Eye and Ear Foundation (Yalong Dang), and an unrestricted fellowship grant from the Third Xiangya Hospital of Central South University (Chao Wang).

## Author contributions statement

Design of the Study (NAL); Conduct of the study (YD, SW, CW, RTL, MS, NAL); Collection, management, analysis, and interpretation of the data (YD, SW, NAL); Preparation and review of manuscript (YD, SW, CW, RTL, MS, NAL). All authors read and approved the final manuscript.

